# Histone acetyltransferases and deacetylases are required for virulence, conidiation, DNA damage repair, and multiple stresses resistance of *Alternaria alternata*

**DOI:** 10.1101/2021.05.14.444270

**Authors:** Haijie Ma, Lei Li, Yunpeng Gai, Xiaoyan Zhang, Yanan Chen, Xiaokang Zhuo, Yingzi Cao, Chen Jiao, Fred G. Gmitter, Hongye Li

## Abstract

Histone acetylation, which is critical for transcriptional regulation and various biological processes in eukaryotes, is a reversible dynamic process regulated by HATs and HDACs. This study determined the function of 6 histone acetyltransferases (HATs) (*Gcn5*, *RTT109*, *Elp3*, *Sas3*, *Sas2*, *Nat3*) and 6 histone deacetylases (HDACs) (*Hos2*, *Rpd3*, *Hda1*, *Hos3*, *Hst2*, *Sir2*) in the phytopathogenic fungus *Alternaria alternata* by analyzing targeted gene deletion mutants. Our data provide evidence that HATs and HDACs are both required for mycelium growth, cell development and pathogenicity as many gene deletion mutants (*ΔGcn5*, *ΔRTT109*, *ΔElp3*, *ΔSas3*, *ΔNat3*, *ΔHos2*, and *ΔRpd3*) displayed reduced growth, conidiation or virulence at varying degrees. In addition, HATs and HDACs are involved in the resistance to multiple stresses such as oxidative stress (*Sas3*, *Gcn5*, *Elp3*, *RTT109*, *Hos2*), osmotic stress (*Sas3*, *Gcn5*, *RTT109*, *Hos2*), cell wall-targeting agents (*Sas3*, *Gcn5*, *Hos2*), and fungicide (*Gcn5*, *Hos2*). *ΔGcn5*, *ΔSas3* and *ΔHos2* displayed severe growth defects on sole carbon source medium suggesting a vital role of HATs and HDACs in carbon source utilization. More SNPs were generated in *ΔGcn5* in comparison to wild-type when they were exposed to ultraviolet ray. Moreover, *ΔRTT109*, *ΔGcn5* and *ΔHos2* showed severe defects in resistance to DNA-damaging agents, indicating the critical role of HATs and HDACs in DNA damage repair. These phenotypes correlated well with the differentially expressed genes in *ΔGcn5* and *ΔHos2* that are essential for carbon sources metabolism, DNA damage repair, ROS detoxification, and asexual development. Furthermore, *Gcn5* is required for the acetylation of H3K4. Overall, our study provides genetic evidence to define the central role of HATs and HDACs in the pathological and biological functions of *A. alternata*.

**Importance:** In filamentous fungi, HATs and HDACs have been found to be involved in growth, development, virulence, synthesis of secondary metabolites and multi-stress resistance. However, the role of all HATs and HDACs in the same fungal pathogen has not been systematically explored. Our study revealed that HATs and HDACs are required for the vegetative growth, conidiation, pathogenicity, multiple stresses resistance, DNA damage repair, and carbon source utilization in *A. alternata*. Moreover, HATs and HDACs identified in *A. alternata* are highly conserved in many notorious pathogenic fungi, such as *Magnaporthe grisea* and *Candida albicans*. Therefore, this study not only systematically reveals the biological functions of twelve histone acetylation-required enzymes for the first time, but also provides a foundation for future investigations in *A. alternata* or other pathogenic fungi.

## Introduction

The fundamental structural subunit of the chromatin is the nucleosome, which consists of approximately 200 bp of DNA wrapped around an octamer of core histone proteins H2A, H2B, H3 and H4 (1). The reversible acetylation of specific lysine residues in N-terminal tails of core histones is maintained by histone acetyltransferases (HATs) and histone deacetylases (HDACs) and has been found to play a pivotal role in chromatin-regulated DNA events, such as DNA replication, damage repair, recombination, and gene expression in eukaryotes (2, 3). HATs are classified into five families, including MYST (Ybf2/Sas3, Sas2, MOZ, Tip60), GNAT (Gcn5-related N-acetyltransferases), p300/CBP, nuclear receptor cofactors and basal transcription factors (4). HDACs are grouped into four groups based on phylogenetic analysis and sequence homology, including Class I and II (Rpd3,Hda1), Class III (Sir2 or Sir2-like protein) and Class IV (HDAC11) (4, 5).

In yeast, many HATs or HDACs have been found to be involved in transcriptional regulation and many cellular functions. Gcn5 belongs to the GNAT family and acts as a core subunit of several histone acetyltransferase (HAT) complexes, such as ADA, SAGA or SALSA (6). Gcn5 acetylates N-terminal lysine (K) residues of histones H2B (K11/16) or H3 (K14/18/23/27/36), and the loss of its function leads to defects in gene transcription, sexual differentiation and multiple stresses resistance (7–9). H3K9/27/56 can be acetylated by RTT109, a HAT of p300/CBP family (10–12). In yeast, H3K56ac regulates expression homeostasis and resistance to DNA-damaging agents (13–16). Both Sas3 and Esa1 belong to MYST family, which function as the subunits of Nucleosome Acetyltransferase of histone H3 (NuA3) and H4 (NuA4) complexes, respectively (3). Studies revealed that Esa1 is essential for cell cycle progression in yeast (17). In *S. cerevisiae*, the transcription of FLO1 was significantly reduced in the absence of both Sas3 and Ada2 (18). Rpd3 is a founding member of class I and comprises an N-terminal deacetylase domain (19). In yeast, Rpd3 functions as a catalytic subunit in sin3 complexes and has been linked to many biological functions, such as transcriptional regulation, DNA repair, replication, and ROS resistance (20–23). Hos2 and Hda1 belong to class II, and both of them also contain an N-terminal deacetylase domain (2). In yeast, Hda1 and Rpd3L contribute to the repair of replication-born DSBs (double strand breaks) by facilitating cohesin loading thus preventing genome instability (24). Hos2 acts as a key subunit of Set3 complex which deacetylates H4 and H3 by counteracting Esa1 (25, 26). The yeast Hos2 is involved in various biological functions, such as these involved in cell integrity pathway, cell resistance to multiple stresses, DNA damage repair and expression of growth-related genes (25, 27–30). The best-characterized member in Class III is the NAD-dependent histone deacetylase (HDAC) Sir2. Yeast Sir2 acts as anti-aging gene required for cellular lifespan regulation by caloric restriction (31, 32).

In filamentous fungi, HATs and HDACs have been found to be involved in growth, development, virulence, synthesis of secondary metabolites and multi-stress resistance. Studies in *Aspergillus nidulans* showed that GcnE (ortholog of Gcn5) controls asexual reproduction, and the biosynthesis of sterigmatocystin, penicillin and terrequinone A (33, 34). Genetic deletion of *Gcn5* leads to severe defects in virulence, sexual or asexual development, and/or virulence in *Ustilago maydis*, *Beauveria bassiana*, and *Trichoderma reesei* (35–37). In *B. bassiana* and *C. albicans*, *RTT109* deletion mutant shows attenuated virulence, and increased sensitivity to the stress of oxidation and DNA-damaging agents (38, 39). Many other HATs also have been found to be involved in cell growth and pathogenicity in *C. albicans* (Esa1, Sas2), *M. oryzae* (Sas3), *F. graminearum* (Elp3), and so on (40–42). HDAC Hos2 acts as a vital subunit of Tig1 complex crucial for conidiation and infectious growth in *M. oryzae*, and is required for pathogenesis and dimorphic switch in *U. maydis* (43, 44). Rpd3/Hda1 family of HDACs regulates azole resistance in *C. albicans* (45). Although many HATs and HDACs have been identified to be critical for an array of biological processes in many fungi, the specific function of each gene is not completely the same among different fungi.

*Alternaria alternata* tangerine pathotype is a disastrous fungal pathogen which causes Citrus Brown Spot on young leaves, shoots and fruits of many citrus cultivars. Previous studies revealed that ROS detoxification-related genes such as *Ap1* and *Skn7* (46, 47) or host-selective ACT-toxin biosynthesis genes such as *ACTT5* and *ACTT6* (48), are critical for *A. alternata* to cause lesions on host tissues. Although numerous studies of HATs or HDACs indicate that many of these genes are involved in ROS detoxification, synthesis of secondary metabolites, and various biological functions in many pathogenic fungi, the roles of these genes in *A. alternata* have not been elucidated. In this study, we identified all the HATs and HDACs orthologs in *A. alternata* and clarified their roles in growth and development, carbon source utilization, multiple stresses resistance, DNA damage repair and virulence using genetic and biological analyses. We also performed a transcriptome analysis to explore the regulatory role of *Gcn5* and *Hos2* in this important citrus pathogen.

## Results

### Identification and genetic deletion of HATs and HDACs in *A. alternata*

Seven HATs and six HDACs were identified combining the prediction results of dbHiMo (http://hme.riceblast.snu.ac.kr/) and hmm analysis, and the families to which these genes belong are also listed in **Table 1**. Nucleotide sequence length of corresponding predicted genes ranged from 654 to 3450 bp, the number of introns contained varied from 0 to 8, and the length of proteins varied from 193 to 1082 aa (**Fig 1A and 1B**). Functional domain prediction of amino acid sequences encoded by these genes revealed that all 13 proteins contain functional domains unique to HAT or HDAC, indicating these gene predictions are highly reliable. Mapping these HATs and HDACs to chromosomes revealed that all 13 genes are localized in the essential chromosomes, whereas no HATs or HDACs were identified in conditionally dispensable chromosomes (CDC) (**Fig S1**). Phylogenetic analysis using the neighbor-joining method revealed that all HATs and HDACs predicted in *A. alternata* were highly similar to their corresponding genes in other fungi (**Fig 1C**).

**Fig 1.**
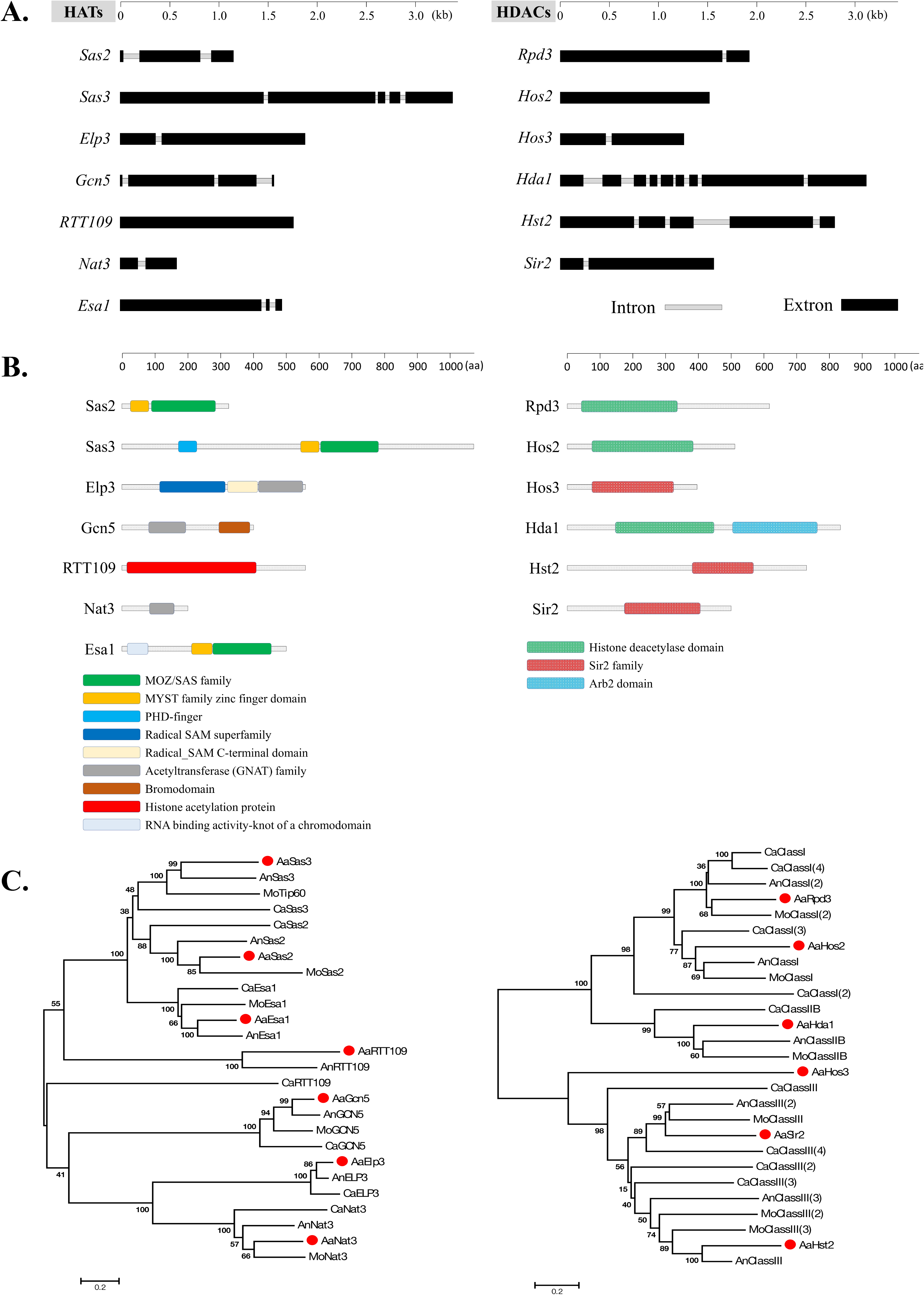
Identification and phylogenetic analysis of HATs and HDACs in *A. alternata*. **(A)** Schematic depiction of ORFs of HATs and HDACs. **(B)** Schematic depiction of conserved domain of HATs and HDACs. **(C)** Phylogenetic trees were constructed by comparing amino acid sequences between different species using the neighbor-joining algorithm of MEGA6. Bar, 5 % sequence divergence. Accession number of amino acid sequences are listed in Table S10.

To investigate the function of HATs and HDACs in *A. alternata*, mutants defective for *Gcn5* (*ΔGcn5*), *Elp3* (*ΔElp3*), *RTT109* (*ΔRTT109*), *Nat3* (*ΔNat3*), *Sas3* (*ΔSas3*), *Sas2* (*ΔSas2*), *Hos2* (*ΔHos2*), *Rpd3* (*ΔRpd3*), *Hos3* (*ΔHos3*), *Hda1* (*ΔHda1*), *Hst2* (*Δ Hst2*) and *Sir2* (*Δ Sir2*) were generated by the homologous recombination method. The mutants were verified by PCR analysis (**Fig S2**). However, no *Esa1* deletion mutants were obtained after multiple attempts.

### HATs and HDACs are required for growth and cell development

In HATs disrupted mutants, vegetative growth rate of *ΔGcn5*, *ΔElp3*, *ΔRTT109*, *ΔNat3* and *ΔSas3* was reduced by 66%, 44%, 12%, 71% and 41% compared to wild-type on potato dextrose agar (PDA), indicating that most HATs play a vital role on the growth of *A. alternata* (**Fig 2A**). In addition, *ΔGcn5* and *ΔNat3* produced fewer aerial hyphae than wild-type on PDA. Microscopic examination revealed that the impacts of different HATs on the development of *A. alternata* varied greatly. The *ΔGcn5* strain produced swollen and stubby hyphae and more hyphae branches, but failed to produce any spores (**Fig 2B, 2C**, **S3**). No spores were observed in *ΔNat3* after 6 days of incubation, but a few spores appeared as incubation time increased. The hyphal tip of *ΔNat3* was irregularly twisted and the hyphae were frequently branched. The conidiation of *ΔElp3*, *ΔSas3* and *ΔRTT109* decreased significantly, but the mycelium morphology remained the same as the wild-type level. Only *ΔSas2* displayed wild-type levels of vegetative growth, hyphal development and conidiation.

**Fig 2.**
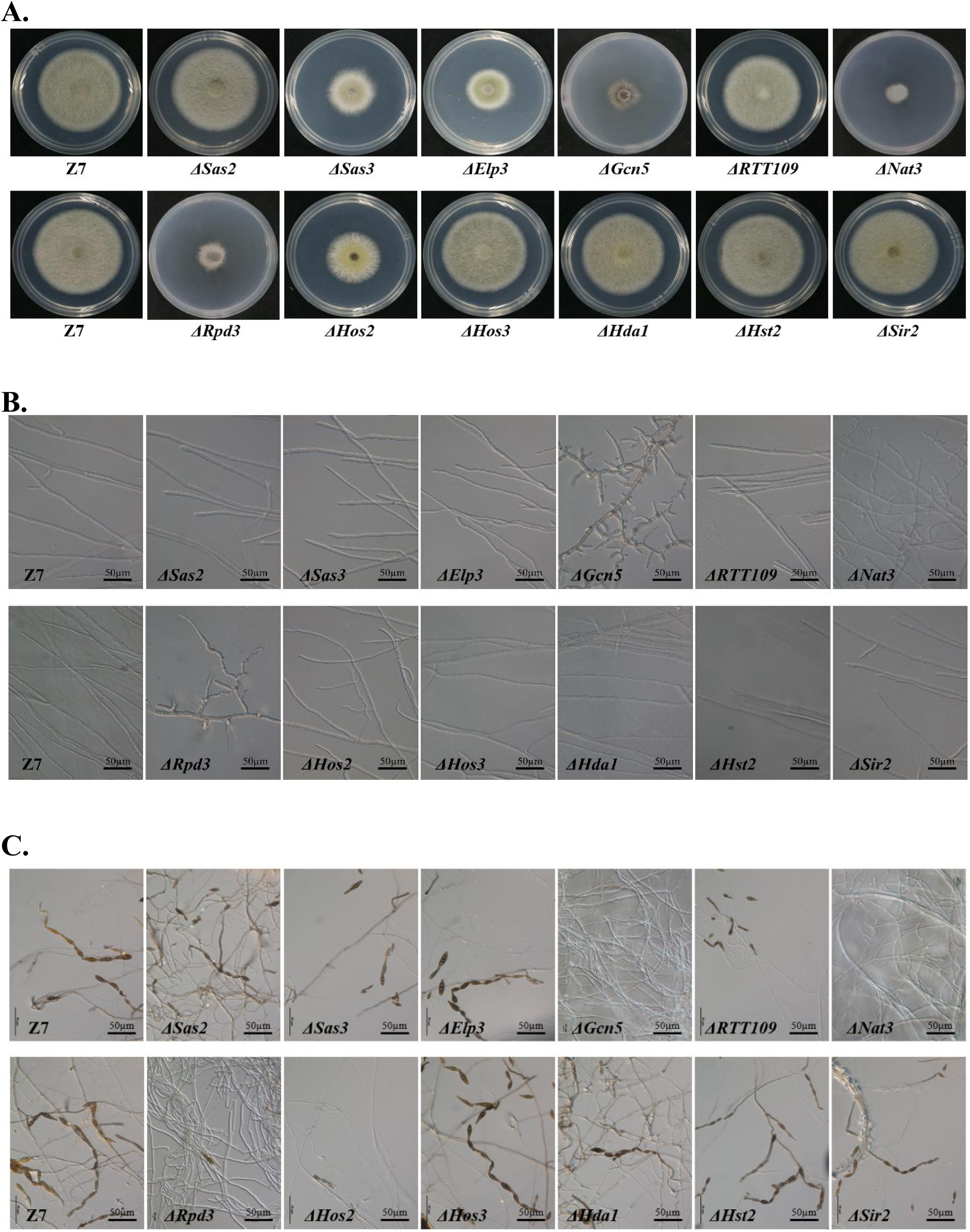
HATs and HDACs were required for growth and development. **(A)** Radial growth of the wild-type strain (Z7) and mutants. **(B)** Conidiation of Z7 and mutants. **(C)** Hyphae morphology of Z7 and mutants.

Unlike HATs, few HDACs were involved in vegetative growth and conidiation. Only *ΔHos2* and *ΔRpd3* grew significantly slower (with inhibition of 37% and 67%, respectively) than the wild-type (**Fig 2A**). Microscopic examination revealed that conidiation was inhibited significantly (>90%) in *ΔHos2* and *ΔRpd3*. Similar to *ΔGcn5*, *ΔRpd3* also produced stubby and swollen hyphae (**Fig 2B**). *ΔHos3*, *ΔHda1*, *ΔHst2*, and *ΔSir2* displayed wild-type levels of vegetative growth and conidiation (**Fig2C**).

### HATs and HDACs are required for fungal virulence

To determine the effect of HATs and HDACs on the pathogenicity of *A. alternata*, virulence assays were performed on detached Dancy leaves. For HATs deletion mutants, *ΔGcn5*, *ΔSas3* and *ΔNat3* failed to induce visible lesion on citrus leaves. *ΔRTT109* and *ΔElp3* induced smaller necrotic lesions than those induced by the wild type 2 days post inoculation (dpi) (**Fig 3**). Only *ΔSas2* induced necrotic lesions at a rate and magnitude comparable to those for the wild-type on Dancy leaves 2 dpi. For HDACs deletion mutants, *ΔRpd3* and *ΔHos2* induced smaller necrotic lesions compared to those induced by the wild type at 2 dpi. *ΔHos3*, *ΔSir2* and *ΔHda1* produced necrotic lesions similar to those induced by Z7 (**Fig 3**). Quantitative analysis revealed that the area of lesion induced by the *ΔRTT109*, *ΔElp3*, *ΔSas2*, *ΔHos2*, *ΔRpd3*, *ΔHda1*, *ΔHos3*, *ΔHst2*, and *ΔSir2* were about 79%, 32%, 92%, 18%, 11%, 97%, 104%, 96%, and 103%, respectively, of those induced by Z7.

**Fig 3.**
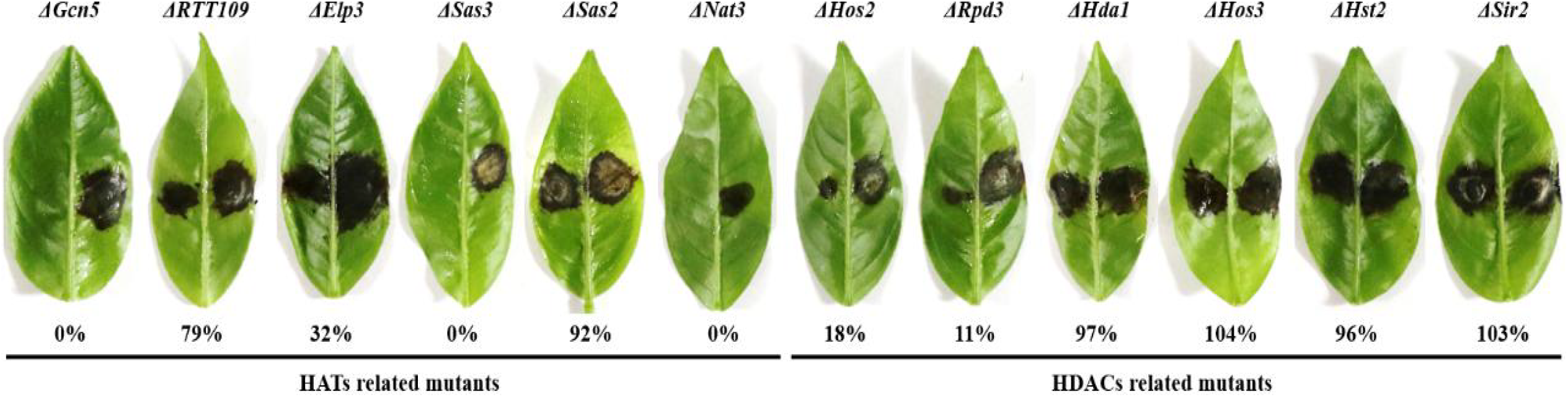
HATs and HDACs were required for *A. alternata* pathogenicity. Pathogenicity was assayed on detached Dancy leaves by placing 5 mm agar plug covering fungal mycelium on the bottom of leaves (left, mutants; right, Z7). The leaves were incubated in a plastic box for lesion development and the necrotic lesions was observed at 2 days post inoculation (dpi). The percent changes of necrotic lesions appearing on the leaves calculated in relation to those induced by Z7 are also indicated.

### HATs and HDACs coordinate carbon sources utilization

To examine whether HATs or HDACs were required for the utilization of carbon sources, mycelial radical growth assays were performed on MM medium supplemented with sucrose, starch, glucose or lactose as sole carbon sources (**Fig 4**). In HATs disruption mutants, the vegetative growth inhibition rate of *ΔGcn5* on all tested sole carbon source media for 4 days was 100%. At the same time, the growth inhibition ratio of *ΔSas3* on the MM medium supplemented with glucose, sucrose, lactose and starch was increased by 28%, 41%, 39% and 41%, compared to the wild type. In contrast, *ΔSas2*, *ΔElp3*, and *ΔRTT109* displayed wild-type sensitivity to all tested sole carbon source media. Most HDACs disruption mutants, including *ΔSir2*, *ΔHos3*, *ΔHda1*, and *ΔHst2*, exhibited wild-type sensitivity to all test medium. Only *ΔHos2* showed slightly increased sensitivity on carbon sources of glucose and starch.

**Fig 4.**
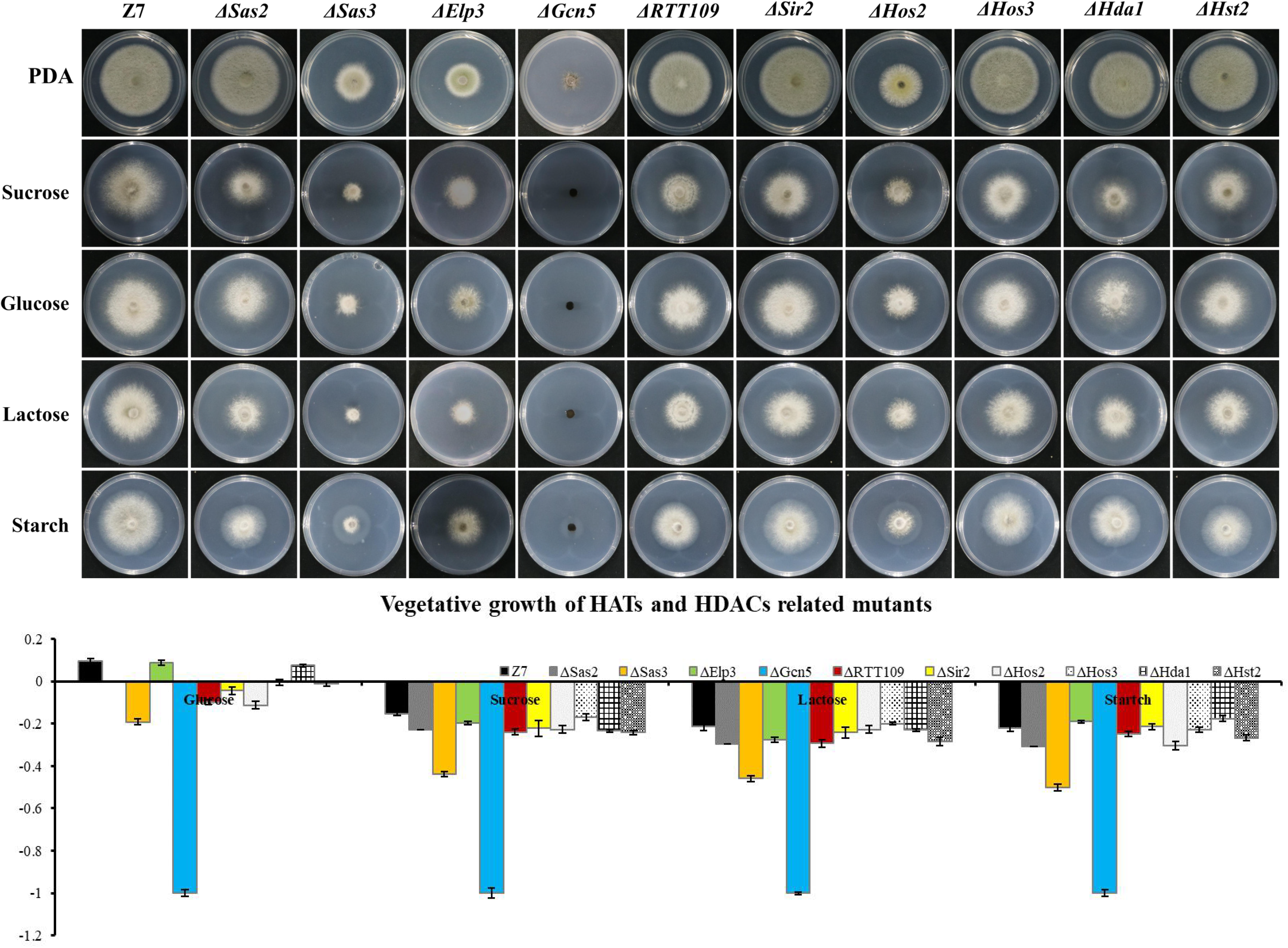
HATs and HDACs were involved in carbon source utilization. Z7, and mutants with impaired HATs or HDACs cultured on MM medium amended with sucrose, glucose, lactose, or starch as sole carbon source. The percentage of growth reduction determined by comparing a cumulative percentage of the growth of Z7 and mutants grown on same medium is also shown.

To prove further that the severe growth defects of *ΔGcn5* on sole carbon source medium was not caused by its slow vegetative growth, the incubation time was extended to 10 days. Growth examination indicated that the growth inhibition ratio of *ΔGcn5* on the medium with lactose as sole carbon source was still 100%, and only a few hyphae of *ΔGcn5* appeared on carbon sources of sucrose, starch or glucose (the growth inhibition ratio is close to 100%), confirming that the *Gcn5* was required for the utilization of carbon sources (**Fig S4**).

### HATs and HDACs are involved in resistance to multiple stresses

In mutants defective for HATs, both *ΔElp3* and *ΔGcn5* were more sensitive to H_2_O_2_ and CHP (cumyl-H_2_O_2_), while *ΔSas3* and *ΔRTT109* just showed increased sensitivity to H_2_O_2_, compared with wild-type. In addition, *ΔSas3* showed elevated sensitivity to cell wall-targeting agents such as Congo red (CR) and sodium dodecyl sulfonate (SDS). At the same time, *ΔElp3*, *ΔGcn5*, and *ΔRTT109* just displayed increased sensitivity to sodium dodecyl sulfonate. *ΔSas3*, *ΔGcn5*, and *ΔRTT109* displayed increased sensitivity to NaCl, KCl and CuSO_4_. Only *ΔRTT109* displayed increased sensitivity to CaCl_2_. However, *ΔSas3*, *ΔElp3*, and *ΔGcn5* displayed enhanced resistance to CaCl_2_ compared to wild-type. Both *ΔElp3* and *ΔGcn5* also showed increased tolerance to V8 medium compared to wild-type. Only *ΔSas2* showed wild-type resistance to all test chemicals (**Fig 5**).

**Fig 5.**
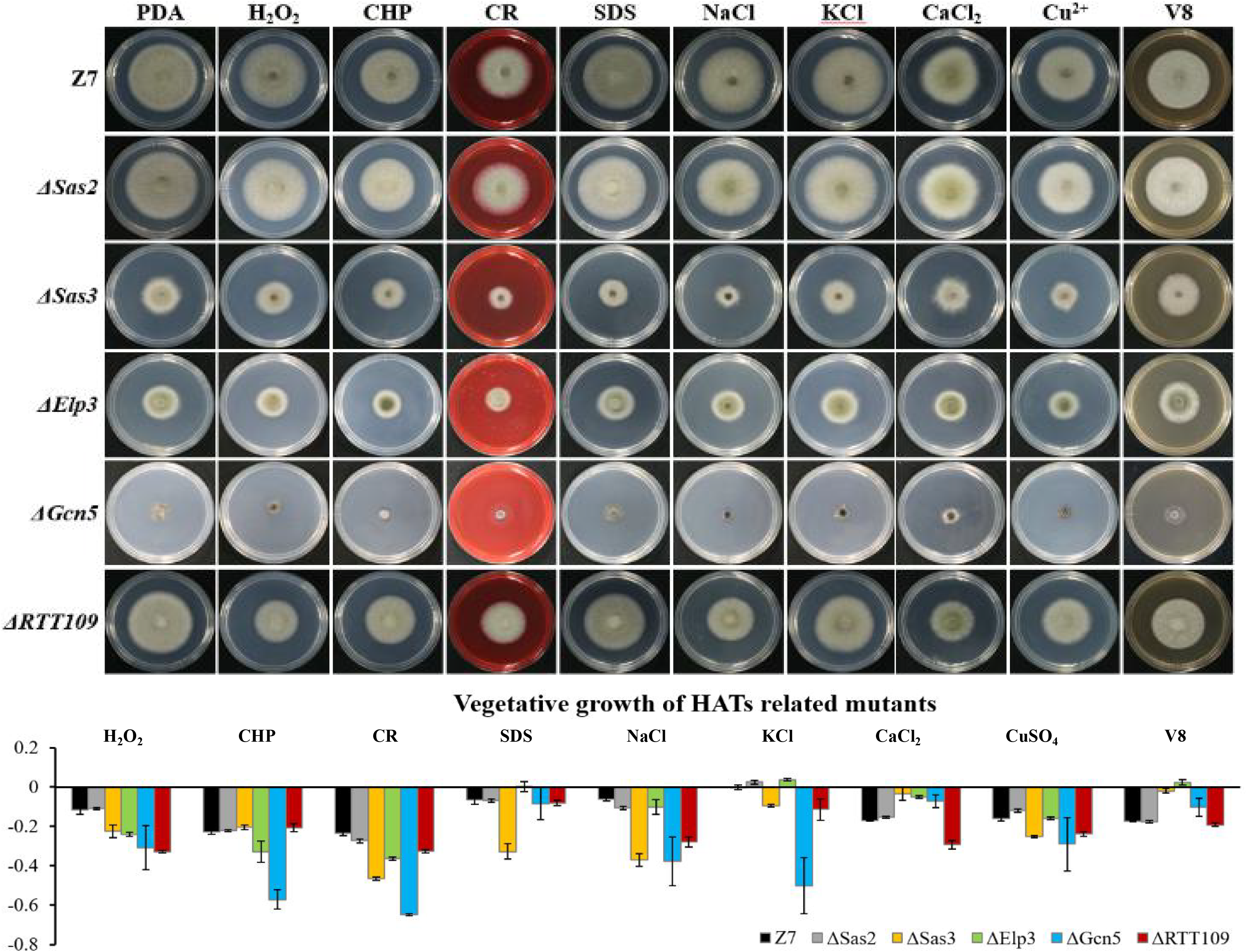
Chemical sensitivity assays. Z7 and HATs related mutants were grown on PDA with or without the indicated chemicals [10 mM H_2_O_2_, 1 mM cumyl hydroperoxide (CHP), 1 M KCl, 1 M NaCl, 250 mM CaCl_2_, 1 mM CuSO_4_, 0.01 % sodium dodecyl sulfate (SDS), 0.2 mg/ml congo red (CR)]. The percentage of growth reduction determined by comparing a cumulative percentage of the growth of Z7 and mutants grown on same medium is also shown.

Unlike HATs which were widely involved in the adaption to various stresses, most of HDACs such as *Hos3*, *Hda1*, *Hst2* and *Sir2* were not involved in the resistance to most tested chemicals. Sensitivity assays revealed that *ΔHos2* showed increased sensitivity to H_2_O_2_, CHP, CR, SDS, NaCl, KCl, CaCl_2_ and V8 medium, but not sorbitol and CuSO_4_ (**Fig 6**).

**Fig 6.**
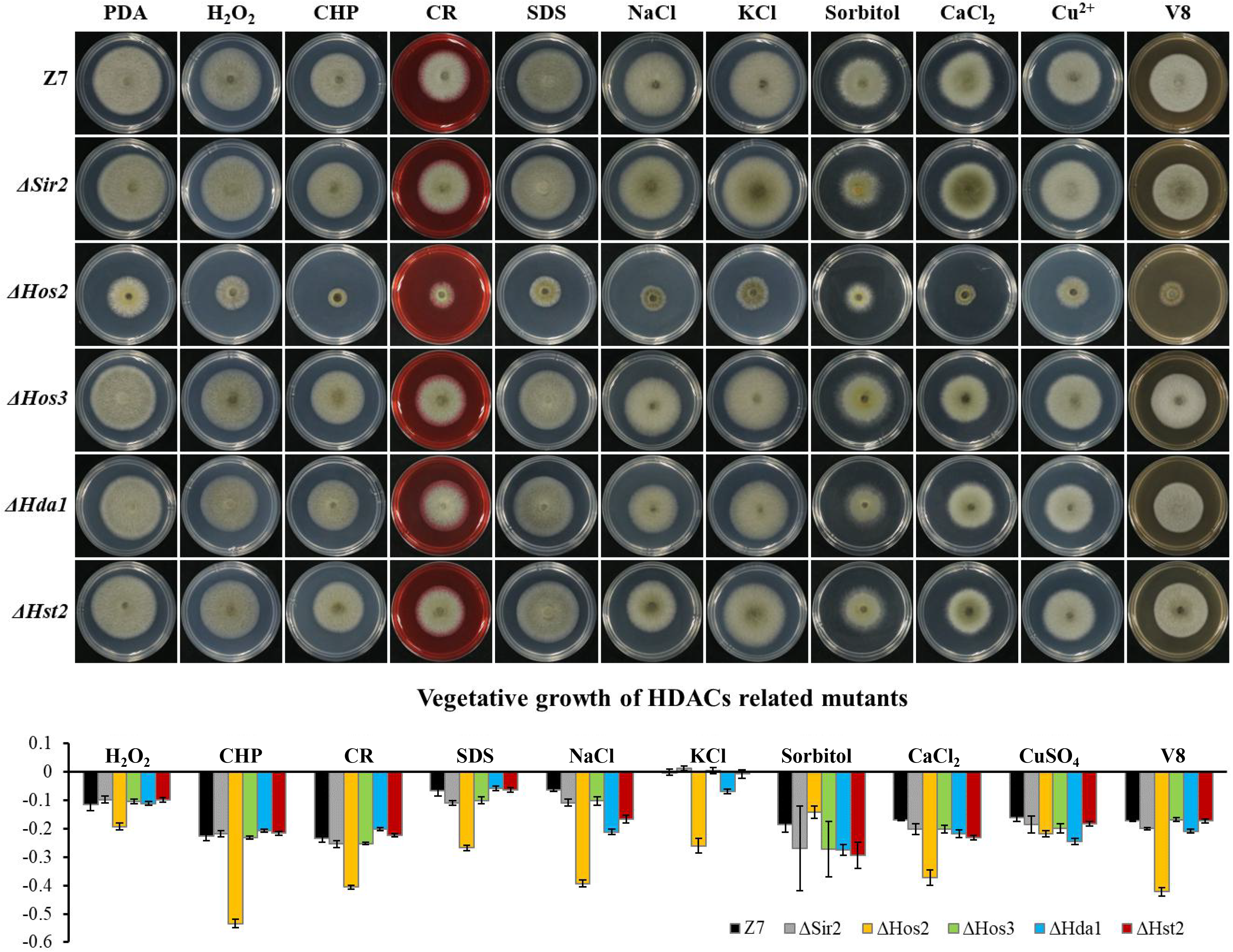
Chemical sensitivity assays. Z7 and HDACs related mutants were grown on PDA with or without the indicated chemicals [10 mM H_2_O_2_, 1 mM cumyl hydroperoxide (CHP), 1 M sorbitol, 1 M KCl, 1 M NaCl, 250 mM CaCl_2_, 1 mM CuSO_4_, 0.01 % sodium dodecyl sulfate (SDS), 0.2 mg/ml congo red (CR)]. The percentage of growth reduction determined by comparing a cumulative percentage of the growth of Z7 and mutants grown on same medium is also shown.

Assays for fungicide sensitivity indicated that both *Δ Gcn5* and *Δ Hos2* displayed increased sensitivity to chlorothalonil, but not mancozeb and boscalid. All the other HATs or HDACs mutants displayed wild-type sensitivity to the test fungicides (**Fig S5**).

### Transcriptome analysis defines the global regulatory role of Gcn5 and Hos2

The results described above indicated that both *Gcn5* and *Hos2* played vital roles in vegetative growth, development, multiple stresses resistance, carbon sources utilization, ROS detoxification, and virulence. To explore the transcriptional regulatory mechanism of *Gcn5* and *Hos2*, transcriptome analyses of *Gcn5* and *Hos2* compared to wild type were performed. Overall, 1503 up-regulated and 1051 down-regulated genes were identified in *ΔGcn5* (**Table S1**). In *ΔHos2*, 469 up-regulated and 536 down-regulated genes were identified (**Table S2**). *Gcn5* and *Hos2* were one of the genes with lowest transcription level in *ΔGcn5* and *ΔHos2*, respectively, indicating the genetic deletion of *Gcn5* and *Hos2* was reliable (**Table S1, S2**). In addition, RT-qPCR results of 14 randomly selected differentially expressed genes (DEGs) was consistent with transcriptome data of *ΔGcn5* and *ΔHos2*, indicating that the transcriptome data were reliable. Although the DEGs in *ΔGcn5* and *ΔHos2* accounted for up to 20% and 8% of all genes in the genome, respectively, the transcription levels of the other 12 HATs and HDACs in *ΔGcn5* or *ΔHos2* did not change significantly, indicating that *Gcn5* and *Hos2* are not required for the transcription of the other HATs and HDACs.

COG (Clusters of Orthologous Groups of proteins) analysis of *ΔGcn5* and *ΔHos2* revealed that many DEGs were enriched in the metabolism category, such as Carbohydrate transport and metabolism (G), Amino acid transport and metabolism (E), Lipid transport and metabolism (I), and Inorganic ion transport and metabolism (P). There are also many DEGs in *ΔGcn5* or *ΔHos2* enriched in Transcription (K) and Replication, recombination and repair (L) (**Fig S6, S7**). KEGG enrichment analysis revealed that the most affected metabolism-related pathways in *ΔGcn5* or *ΔHos2* were mainly related to carbon and nitrogen sources metabolism, such as starch and sucrose metabolism, galactose metabolism, Fructose and mannose metabolism, Methane metabolism, and Nitrogen metabolism. In addition, many DEGs in *ΔGcn5* or *ΔHos2* were enriched in genetic information processing, such as DNA damage repair, RNA transport, and tRNA biosynthesis (**Fig S6, S7**). Although many DEGs in *ΔGcn5* and *ΔHos2* were enriched in the same pathways, the amount and expression level of those genes varied greatly.

*Gcn5* and *Hos2* are involved in regulating the transcription of many genes involved in carbon (sugar) source utilization. Analysis of DEGs in Glycolysis/Gluconeogensis (ko00010), Fructose and Mannose Metabolism (ko00051), Galactose Metabolism (ko00052), Starch and Sucrose metabolism (ko00500), Pentose Phosphate Pathway (ko00030), Citrate cycle (ko00020) and Pentose and Glucuronate interconversions (ko00040) in *ΔGcn5* and *ΔHos2* revealed that the number of down-regulated genes (*ΔGcn5*, 22; *ΔHos2*, 15) was significantly greater than the up-regulated genes (*ΔGcn5*, 22; *ΔHos2*, 15) (**Fig 7**). In yeast, invertase and hexokinase are critical for the utilization of sucrose, lactose, glucose or fructose (49). In *A. alternata*, AALT_g6039 encodes hexokinase, and AALT_g5797, AALT_g10082 and AALT_g2401 encode invertases. All four genes were significantly down-regulated in *ΔHos2*, but only AALT_g6039 and AALT_g2401 were significantly down-regulated in *ΔGcn5*.

**Fig 7.**
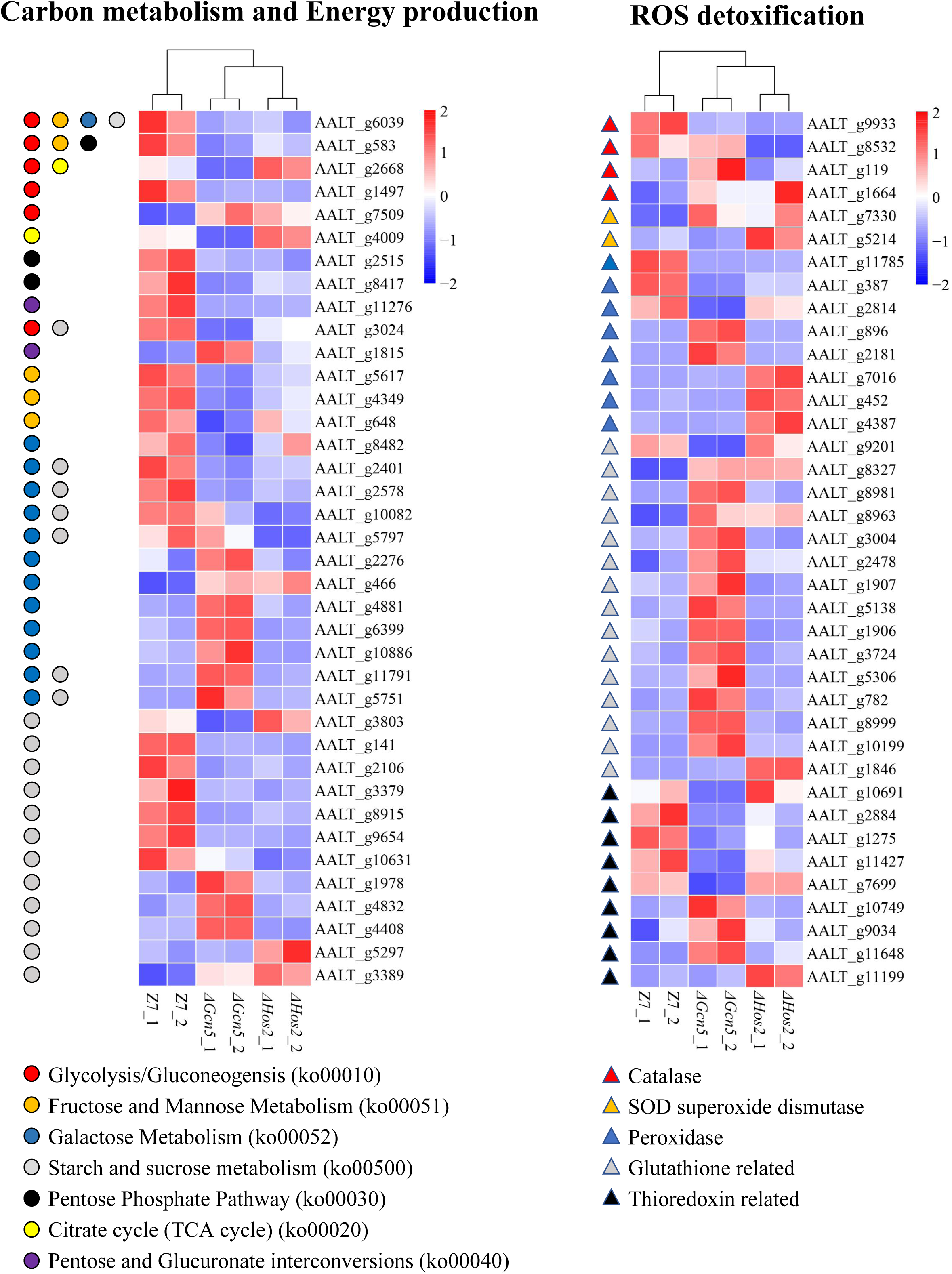
Heatmap of differentially expressed genes enriched in carbon metabolism and energy production, and ROS detoxification.

*Gcn5* and *Hos2* affect the expression of ROS detoxification genes. In view of the increased sensitivity of *ΔGcn5* and *ΔHos2* to oxidative stress, we analyzed the transcription level of 96 ROS detoxification genes, including 7 catalases, 7 SOD superoxide dismutases, 42 glutathione system-related genes, 20 thioredoxin system-related genes and 20 peroxidases (**Table S3**). Although *ΔGcn5* and *ΔHos2* were more sensitive to oxidants, most of DEGs in these 96 ROS detoxification genes were up-regulated. In *ΔGcn5*, 21 DEGs were up-regulated and 10 DEGs were down-regulated (**Fig 7**). In *ΔHos2*, 9 DEGs were up-regulated and 4 DEGs were down-regulated.

*Gcn5* and *Hos2* mediate the expression of genes required for DNA damage repair. Base excision repair (BER), Nucleotide excision repair (NER), Mismatch repair (MMR), Homologous recombination (HR), and Non-homologous end-joining (NHEJ) play important roles in the repair of damaged DNA in organisms. Transcriptome analysis revealed that BER, NER, MMR or HR-related genes including UDG (Uracil-DNA glycosylase, AALT_g5833), Pol δ (DNA polymerase delta, AALT_g8966), and TFIIH (transcription factor b, AALT_g7297) were down-regulated in *ΔGcn5* (**Fig S7, Table S1, S2**). In *ΔHos2*, BER and NER-related gene PARP [poly (ADP-ribose) polymerase, AALT_g9674], and NER related gene XPA (DNA-repair protein, AALT_g6731) were down-regulated (**Table S2**). In addition, AALT_g76 (BER, endonuclease), AALT_g10858 (NER, RING-box protein 1) in *ΔGcn5*, and AALT_g15 (NHEJ, DNA ligase 4) in *ΔHos2* were up-regulated (**Table S2**).

*Gcn5* and *Hos2* regulate expression of genes involved in protein processing and degradation. In *ΔGcn5,* 7 DEGs enriched in “protein processing in endoplasmic reticulum (ko04141)” were up-regulated, including protein glycosyltransferase (AALT_g5706), Mannosyl-oligosaccharide glucosidase (AALT_g11241), Mannosyl-oligosaccharide alpha-1,2-mannosidase (AALT_g2989), HSP20 (AALT_g9536), UBC5 (AALT_g6888), UBC7 (AALT_g619), and RBX1 (AALT_g4961), which are involved in protein processing and degradation. Furthermore, eight up-regulated genes were enriched in “Ubiquitin mediated proteolysis (ko4120)” and “Regulation of Autophagy (ko04140)” (**Fig S8, Table S1**). Only 4 genes were down-regulated in the above three pathways. In *ΔHos2*, all four DEGs enriched in “protein processing in endoplasmic reticulum” (AALT_g173 and AALT_g6960), “Ubiquitin mediated proteolysis” (AALT_g6960), and “Regulation of Autophagy” were up-regulated (AALT_g11308) (**Table S2**).

In addition, deletion of *Gcn5* or *Hos2* also affected the transcription of genes involved in Carbohydrate-active enzymes (CAZymes), secondary metabolite cluster, CYP450, and conidiation (**Table S4, S5, S6, S7, Fig S9**). The transcription of all ACT toxin synthesis-related genes did not change significantly in *ΔGcn5* and *ΔHos2* compared to wild-type. However, AALT_g11758 (CYP450), located in the ACT toxin gene cluster and required for pathogenicity but plays no role for ACT toxin synthesis (50), was significantly down-regulated in both *ΔGcn5* (FC = −25) and *ΔHos2* (FC = −6).

### HATs and HDACs are required for the resistance to DNA-damaging agents

Previous studies showed that *Gcn5*, *Hos2*, and *RTT109* are required for DNA damage repair in *B. bassiana* and *S. cerevisiae* (10, 38, 51). Therefore, we tested whether *RTT109*, *Gcn5* and *Hos2* were involved in the resistance to DNA-damaging agents such as camptothecin (CPT), hydroxyurea (HU) and methyl methyl methanesulfonate (MMS) in *A. alternata*. Compared with wild-type, growth of *Δ RTT109* was significantly suppressed by 40% and 89% on PDA containing 5 μM CPT or 0.1% MMS, respectively (**Fig 8**). However, *ΔRTT109* displayed wild-type radial growth on PDA amended with 5 mM HU. Unlike *ΔRTT109*, the growth of *ΔHos2* on HU-added PDA was inhibited by 20% compared to wild-type. At the same time, *ΔHos2* showed wild-type resistance to 5 μM CPT or 0.1% MMS. In addition, *ΔGcn5* displayed increased sensitivity to 5 mM HU and 0.1% MMS (the growth inhibition rate is 29% and 19%, respectively, compared to wild-type) but displayed wild-type resistance to 5 μM CPT (**Fig 8**).

**Fig 8.**
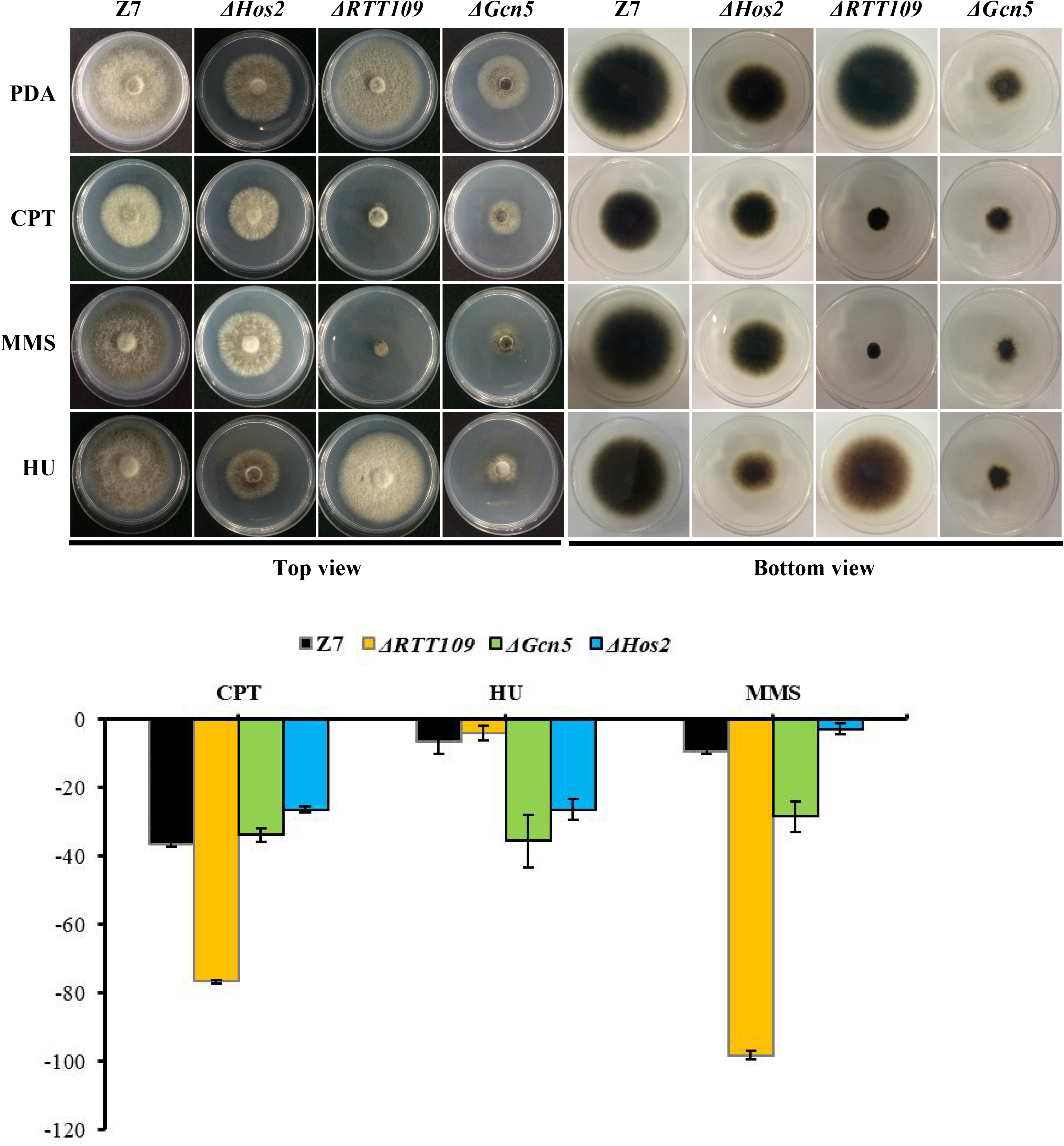
HATs and HDACs were involved in the resistance to DNA-damaging agents. Z7, *ΔHos2*, *ΔRTT109*, and *ΔGcn5* were grown on PDA with or without the indicated chemicals [5 mM hydroxyurea (HU), 0.1 % methyl methanesulfonate (MMS), or 5 μM camptothecin (CPT)]. The percentage of growth reduction determined by comparing a cumulative percentage of the growth of Z7 and mutants grown on same medium is also shown.

### Gcn5 is required for DNA damage repair

In order to analyze the DNA damage repair ability of *Gcn5*, we performed genome resequencing using ultraviolet ray (UV) irradiated *ΔGcn5* and wild-type. In the *Gcn5* coding region, no reads were detected in *ΔGcn5*, but many reads were found in wild-type, which proves once again that the knock-out experiment of *Gcn5* was successful (**Fig 9B**). In *ΔGcn5*, 949 unique SNPs were detected after UV irradiation. However, in UV irradiated wild-type, only 298 unique SNPs were found, which accounted for 31% of *ΔGcn5*, indicating that *Gcn5* is involved in the repair of DNA damage caused by UV (**Fig 9A**).

**Fig 9.**
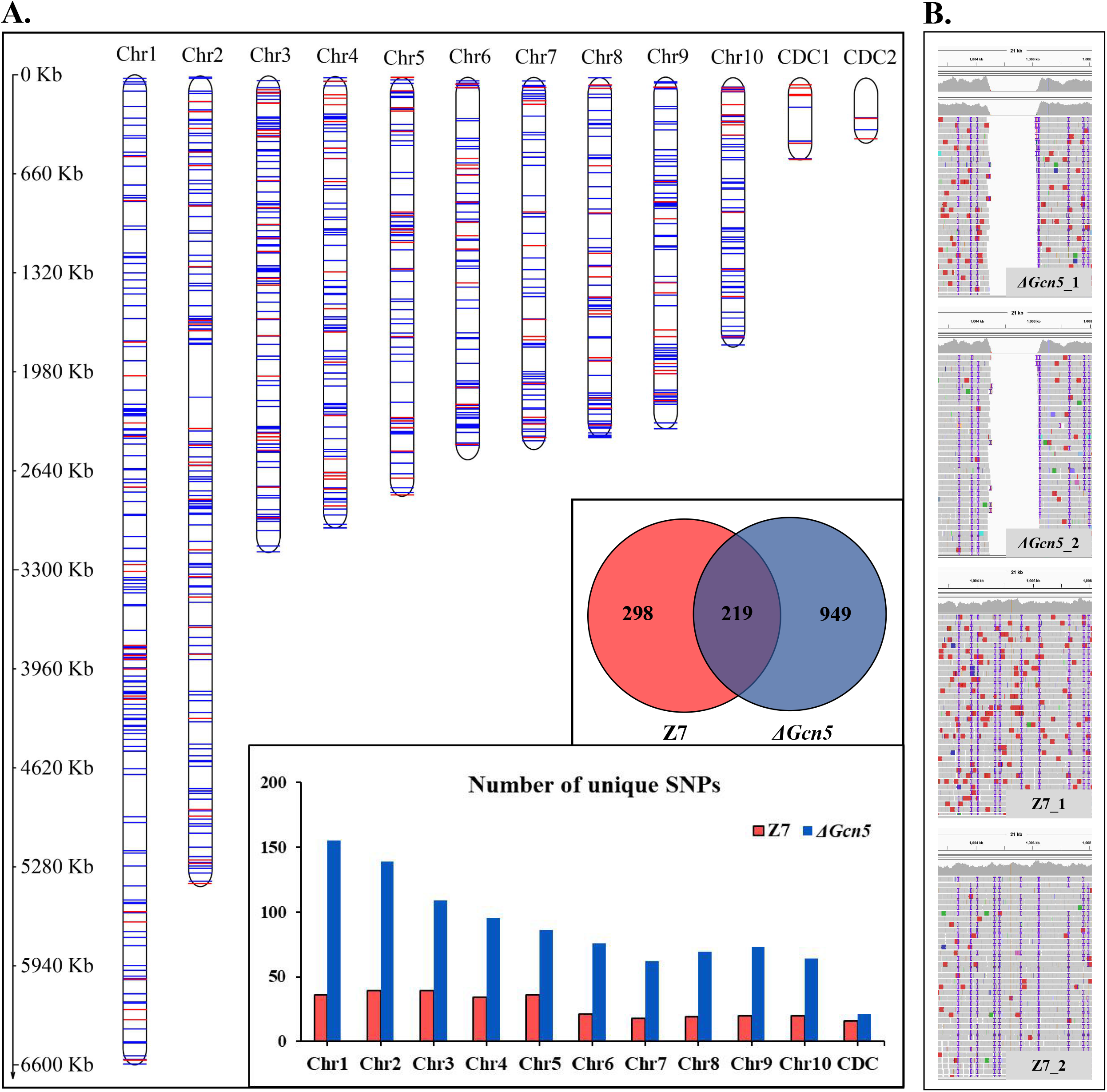
SNPs analysis of *ΔGcn5* and wild-type irradiated by UV. *ΔGcn5* and Z7 grown on PDA were irradiated with UV for 10 seconds every 24 hours, and repeated 4 times. Hyphae were taken using sterilized toothpicks and incubated in PDB (150 rpm, 2 days). DNA was extracted from mycelium collected from PDB for whole-genome resequencing. The protocol for SNPs analysis is described in Experimental procedures.

### Histone acetylation targets of Gcn5

To test whether histone acetylation levels were altered in *ΔSas3* and *ΔGcn5*, western blot was carried out using specific antibodies, directed against H3K4ac, while antibody against H3 was used as a loading control. As shown in **Fig 10**, the level of H3K4ac was significantly decreased (*P* < 0.01) in the *ΔGcn5* compared to Z7. No reduction of the signal for H3K4ac was observed in *ΔSas3*.

**Fig 10.**
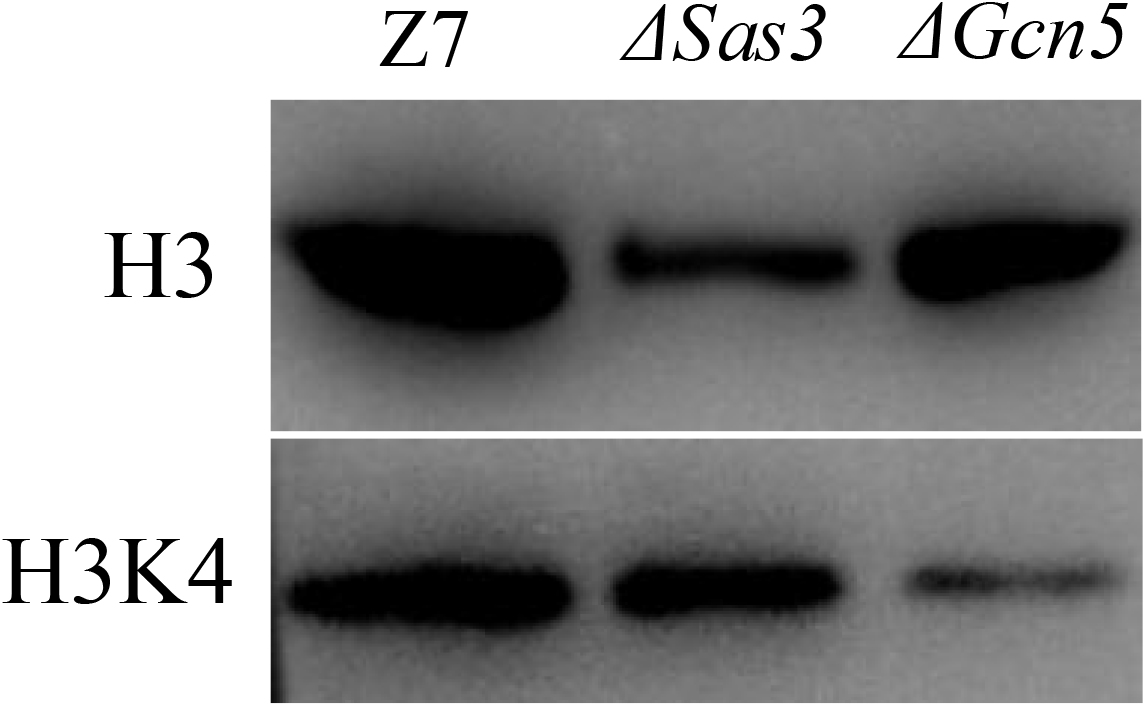
Western blotting analysis of proteins extracted from Z7, *ΔSas3*, and *ΔGcn5*, respectively. **(A)** The respective strains were grown for 36 hours in PDB. The anti-acetyl H2K4 (H3K4ac) and anti-acetyl H3K18 (H3K18ac) were used for detecting alteration of acetylation levels. Antibody H3 was used as a loading reference. (B) Quantification of western blot signals in the three replicates. Data were expressed as mean IOD (relative to H3) ± standard error. \**P* < 0.05, \*\**P* < 0.01.

## Discussion

Histone acetylation, which is important for multiple cellular processes, occurs in different organisms ranging from fungi to mammals, and the acetylation level manifested by HATs and HDACs dynamically changes over time during development and differentiation (3, 52). In plant pathogenic fungi, several HATs and HDACs have been found to be required for growth, virulence and adaptation to environmental stress (41, 52, 53). Although HATs and HDACs are conserved in cells, the functions of these genes are various in different species. In this study, a total of 13 HATs and HDACs were identified in *A. alternata*. To systematically elucidate the functions of HATs and HDACs in *A. alternata*, we created 6 HATs and 6 HDACs deletion mutants. Our studies showed that HATs and HDACs were widely involved in biological processes of mycelial growth and development, conidial production, multiple stresses resistance, carbon source utilization, DNA damage repair, and pathogenicity in *A. alternata*.

As shown in the present study, HATs and HDACs are involved in developmental processes because *ΔGcn5*, *ΔHos2*, *ΔRpd3*, *ΔNat3*, *ΔSas3* and *ΔRTT109* produce no or fewer conidia. In addition, *ΔGcn5*, *ΔRpd3* and *ΔNat2* produces more hyphal branches than wild-type. In some filamentous fungi like *N. crassa* and *A. nidulans*, the central development pathway relies upon the key activator genes *BrlA*, *AbaA* and *WetA* (54, 55). Six upstream developmental activators (*FlbA*, *FlbB*, *FlbC*, *FlbD*, *FlbE*, and *FluG*) are required for the activation of *BrlA* and the initiation of conidiation (56, 57). Our transcriptome data revealed that *FlbA*, *FlbC*, and *FlbD* were down-regulated in *ΔGcn5* (**Table S6**). Previous studies have revealed the conidiation of *A. alternata* is closely regulated by the calcium-mediated signaling pathway, Fus3 and Slt2 MAPK signaling pathway, as well as the cAMP dependent protein kinase A (PKA) (58–61). Other factors including NADPH oxidase (Nox), thioredoxin reductase (Trr1), glutathione reductase (Glr1), nascent polypeptide associated complex α subunit (Nac1), and skn7 response regulator are also required for the developmental processes in *A. alternata* (47, 62–64). Whether or not those regulators interact with HATs or HDACs remains to be determined. In addition to conidia formation, HATs and HDACs are required for vegetative growth. *ΔGcn5*, *ΔNat3*, *ΔSas3*, *ΔRTT109*, *ΔElp3*, *ΔHos2* and *ΔRpd3* displayed growth reduction compared with wild-type, consistent with the findings in *B. bassiana* (*Gcn5*, *RTT109*, *Hos2*, and *Rpd3*), *F. graminearum* (*Gcn5*, *RTT109*, *Elp3*, and *Sas3*), and *S. cerevisiae* (*Nat3*) (35, 38, 42, 51, 52, 65, 66).

Experiments further demonstrated that *ΔGcn5*, *ΔSas3*, *ΔRTT109*, and *ΔHos2* are required for cellular resistance to osmotic stress. In yeast, SAGA (Spt-Ada-Gcn5) is required for the transcription of Hog1-mediated genes under severe osmostress (67). Chip-on-chip experiments revealed that acetylation at lysines 9 and 12 of histone H3 increases in osmostress up-regulated genes and decreases in repressed genes (68). The *Hog1* deletion mutant in *A. alternata* displayed reduced vegetative growth rate and increased sensitivity to KCl and NaCl, which is consistent with the phenotype of *ΔGcn5*, *ΔSas3*, *ΔRTT109* and *ΔHos2*, indicating that transcription of some genes regulated by *Hog1*, *Gcn5*, *Sas3* and *Hos2* may be similar (69). Similar results were observed in *F. graminearum*, in which the *Hog1*, *Gcn5*, *Sas3*, *RTT109* are also involved in vegetative growth and resistance to osmotic stress (52, 70). In addition, *ΔSas3* and *ΔHos2* are more sensitive to cell wall-disturbing compounds including CR and SDS, indicating both of these genes play a positive role in cell wall assembly.

Carbon source (sugar) metabolism, which is one of the most important biological processes in organisms, is involved in growth, development and multiple stress resistance (71, 72). This study found that *Sas3*, *Hos2*, and especially *Gcn5* displayed severe growth defects on sole carbon source medium, suggesting an involvement of these genes in carbon metabolism in *A. alternata*. In *C. albicans*, acetyl-CoA and acetate metabolism play a central role in growth on both glucose and non-glucose carbon sources (73). In *B. bassiana*, deletion of *Rpd3* and *Gcn5* resulted in growth defects on medium modified with different carbon sources, confirming the important role of histone acetylation in carbon source utilization (35, 66). In addition, most DEGs enriched in carbon source metabolism pathways, such as “Glycolysis/Gluconeogensis”, Fructose and Mannose Metabolism”, “Galactose Metabolism”, “Starch and Sucrose Metabolism”, etc., are down-regulated in *ΔGcn5* and *ΔHos2*, making it not surprising to see the reduced carbon source utilization ability in *ΔGcn5* and *ΔHos2*. In microorganisms, glucose metabolites would lead to carbon catabolite repression (CCR) and inhibit the utilization of other carbon sources (74, 75). Sucrose non-fermenting 1 (SNF1) protein kinase complex is required for the utilization of non-fermentable carbon sources in fungi and enable them to adapt to the adversity of glucose deficiency (76). In *Fusarium virguliforme*, the growth of *FvSnf1* deletion mutants on medium amended with galactose as sole carbon source is completely inhibited (77). SNF1 complex-related genes (*AaSnf1*, *AaSip2*, and *AaSnf4*) were not significantly differentially expressed in *ΔGcn5* and *ΔHos2* compared to wild-type. However, the transcription of *SUC2* (AALT_g2401), which is regulated by *Snf1* and is essential for sucrose utilization in *S. cerevisiae* (78, 79), was down-regulated significantly in *ΔGcn5* and *ΔHos2*. Previous studies have shown that Snf1, which regulates Gcn5 occupancy and H3 acetylation at a specific sequence, works in concert with Gcn5 to regulate transcription (80, 81). In addition, similar with *Gcn5* and *Hos2*, *Snf1* also plays an important role in the carbon source utilization in *A. alternata* (63). Therefore, we speculate that the interaction relationship between *Snf1* and these two genes may also exist.

Mutants lacking *Gcn5*, *Sas3*, *Elp3*, *RTT109* or *Hos2* in *A. alternata* were defective in ROS detoxification at varying degrees, indicating all of these five genes are involved in resistance to oxidative stress. *RTT109*, *Gcn5* and *Hos2* in *B. bassiana*, *Elp3* and *Sas3* in *F. graminearum* also play important roles in the resistance to oxidative stress (35, 38, 42, 51, 52). However, *Gcn5* in *Aspergillus flavus* is not required for ROS resistance (82). In addition, *Hos2* plays a negative role in H_2_O_2_ resistance in *M. oryzae*, indicating that the function of specific genes in HATs and HDACs have been differentiated in fungi (83). In *A. alternata*, the ROS detoxification ability is critical for pathogenesis and survival (62, 84, 85). *A. alternata* strains lacking *Nox*, *Hog1*, *Yap1*, *Skn7*, *Gpx3*, or *Nac1* all exhibit increased cellular sensitivity to ROS and decreased pathogenicity on citrus leaves (47, 63, 69, 86–88). Virulence assays revealed that the virulence of *ΔGcn5* and *ΔSas3* was completely lost, which is consistent with the findings in F. *graminearum* (52). Furthermore, the virulence of *ΔElp3*, *ΔRTT109*, and *ΔHos2* decreased significantly, supporting the requirement of ROS detoxification for successful infection by *A. alternata*. In addition, we observed that *Δ Nat3* and *Δ Rpd3* were significantly reduced in pathogenicity and vegetative growth rate. The severe growth defects of *ΔNat3* and *ΔRpd3* may contribute to its reduced virulence. Recent research shows that chlorothalonil causes a redox state change leading to oxidative stress generation in *Danio rerio* (89). Interestingly, sensitivity assays revealed that *A. alternata* strains with impaired *Gcn5* or *Hos2* displayed high sensitivity to chlorothalonil fungicide. In addition, transcriptome data of *ΔGcn5* revealed that all DEGs in the sterol synthesis pathway were significantly down-regulated, including CYP51 homologous genes, which is closely related to DMI fungicide resistance. In *C. albicans*, the loss of *Hda1* and *Rpd3* both caused the pathogen to be significantly more sensitive to azole fungicides, indicating that histone acetylation levels are crucial for pathogens to resist DMI fungicides (45). Transcriptome analysis also revealed that most DEGs involved in ROS detoxification were up-regulated in *ΔGcn5* and *ΔHos2* even without external oxidative stress, implying that the redox state in cells of these two mutants are changed. In eukaryotic cells, endoplasmic reticulum (ER) is essential for folding and trafficking of proteins that enter the secretory pathway, while the disorder of redox homeostasis of ER could result in protein misfolding (90). In *ΔGcn5* and *ΔHos2*, many genes involved in “Protein processing in endoplasmic reticulum” , “Ubiquitin mediated proteolysis” and “Regulation of autophagy” pathways are up-regulated, which mainly play roles in protein folding and degradation.

It is now clear that a subset of HATs and HDACs regulate the genome stability (91, 92). In *B. bassiana*, mutants lacking HATs (*Mst2* and *RTT109*) or HDACs (*Hos2*) displayed increased sensitivity to DNA-damaging stresses, indicating that all of these genes are involved in DNA damage repair (38, 51, 93). In our study, many DEGs are enriched in DNA damage repair pathways. Moreover, *ΔGcn5*, *ΔRTT109*, and *ΔHos2* displayed hypersensitivity to a variety of DNA-damaging agents such as MMS, HU, or CPT, all of which induce replication fork collapse or stalling (94–96), suggesting *Gcn5*, *RTT109* and *Hos2* play an important role in DNA damage repair in *A. alternata*. This speculation was evidenced by the result that more SNPs were generated in *ΔGcn5* in comparison to wild-type when both of them were irradiated by UV.

In summary, our study revealed that HATs and HDACs are required for the vegetative growth, conidiation, pathogenicity, multiple stresses resistance, DNA damage repair, and carbon source utilization in *A. alternata*. In addition, *Gcn5* and *Hos2* play direct or indirect roles in the transcriptional regulation of genes involved in carbon source metabolism, DNA damage repair, ROS detoxification, protein processing and degrading, ergosterol synthesis, and secondary metabolite synthesis.

## Experimental procedures

### Fungal strains and culture conditions

The wild-type Z7 strain of *A. alternata* (Fr.) Keissler used in the mutagenesis experiments was isolated from an infected citrus (*Citrus suavissima* Hort. Ex Tanaka) in Zhejiang, China (97, 98). Fungal strains were cultured on potato dextrose agar (PDA) at 26°C and conidia were collected after incubating on V8 medium for 8 days. Mycelium cultured in liquid potato dextrose broth (PDB) was collected by passing through cheesecloth and used for purification of DNA or RNA.

### Targeted gene disruption and genetic complementation

*ΔSas2*, *ΔSas3*, *ΔElp3*, *ΔGcn5*, *ΔRTT109*, *ΔNat3*, *ΔRpd3*, *ΔHos2*, *ΔHos3*, *ΔHda1*, *ΔHst2* and *ΔSir2* strains were created by deleting *Sas2*, *Sas3*, *Elp3*, *Gcn5*, *RTT109*, *Nat3*, *Rpd3*, *Hos2*, *Hos3*, *Hda1*, *Hst2* and *Sir2* respectively by integrating a bacterial *HYG* cassette under control of *TrpC* gene promoter and terminator in the genome of Z7 using a split marker approach mediated by protoplasts transformation as described previously (62, 99). Fungal transformants were recovered from PDA containing 100 μg/ml hygromycin and examined by PCR with primers specific to its targeted gene. Oligonucleotide primers used in this study are listed in **Supplementary Table S9**.

### Microscopy

Morphological observation of conidia and hyphae was carried using a Nikon microscope equipped with a LV100ND image system (Nikon, Japan).

### Virulence Assays

Fungal virulence was assessed on detached Dancy (*Citrus reticulata* Blanco) leaves inoculated by placing a 5-mm dia. agar plug covered with fungal mycelium on each spot, and those inoculated leaves were kept in a plastic box at 26 °C for 2 to 4 days for lesion development.

### Phenotypic experiments

All the phenotypes of mutants and wild-type were evaluated. To examine the vegetative growth under multiple stresses, mutants and wild-type were inoculated on PDA containing 10 mM H_2_O_2_, 1 mM cumyl hydroperoxide (CHP), 1 M sorbitol, 1 M KCl, 1 M NaCl, 250 mM CaCl_2_, 1 mM CuSO_4_, 0.01 % sodium dodecyl sulfate (SDS), 0.2 mg/ml congo red (CR), 10 mg/L chlorothalonil, 2 mg/L Mancozeb, 3 mg/L Boscalid, 5 mM hydroxyurea (HU), 0.1 % methyl methanesulfonate (MMS), or 5 μM camptothecin (CPT). To examine the carbon utilization ability, mutants and wild-type were inoculated on MM medium using 20 g/L glucose, 10 g/L sucrose, 10 g/L starch, or 10 g/L lactose as sole carbon source. All tests were repeated at least twice with three replicates of each treatment.

### Transcriptome analysis

Wild-type, *ΔGcn5* and *ΔHos2* each with two replicates were cultured in PDB (120 rpm, 28°C) and were harvested after 36 h. RNA was extracted using Axygen RNA purification kit (Capital Scientific, Union City, CA, United States). The libraries were produced using an IlluminaTruSeq RNA Sample Preparation Kit, and they were sequenced on an Illumina HIseq 2500 platform, generating 150 bp paired-end reads. Trimmomatic Ver 0.36 (100) was used to remove adaptors and low-quality reads. TopHat2 (101) was used to map sequences to the reference genome and the mapped reads on each gene were determined by HTSeq (102). Differential expression analysis was determined using DESEQ2 (103) based on the overall transcript counts after normalization. Transcripts with an adjusted FDR less than 0.01 and the absolute value of log2FC (log2 fold change) greater than 2 were considered to be differentially expressed genes (DEGs). DEGs were annotated by searching against the NCBI nr databases. GO and KEGG pathway were performed using clusterProfiler v3.6 (104). anti-SMASH 4.0 (105) was used to predict gene clusters associated with secondary metabolites. Protein domains were predicted in SMART database available at http://smart.embl-heidelberg.de/. Carbohydrate-active enzymes (CAZymes) were predicted using dbCAN meta server (http://bcb.unl.edu/dbCAN2/blast.php). The transcriptomes raw data (**SUB7740259**) have been deposited in NCBI’s Sequence Read Archive.

### Whole-genome resequencing analysis

A paired-end library of each sample was constructed and sequenced by Hiseq2500, with insert sizes of approximately 350bp. Trimmomatic was used to remove adaptors and low-quality reads. The high-quality resequencing reads was mapped to the *Alternaria alternata* reference genome with Bowtie2 using the default parameters. The aligned results were further filtered for unmapped reads and duplicated reads using SAMtools v1.4 and Picard packages, respectively. SNP calling was carried out using Freebayes. Genomic variation analysis was performed using VCFtools. IGV was used to visualize reads on the genome. MapChart was used to visualize SNPs on the genome. The raw data of whole-genome resequencing (**SUB7740329**) have been deposited in NCBI’s Sequence Read Archive.

### Gene expression analyses

Quantitative Real-time PCR (qRT-PCR) was carried out on a 7300 Real Time PCR system (Applied Biosystems, Carlsbad, CA, United States) to validate the transcriptome data. RNA was extracted with an Axygen RNA purification kit (Capital Scientific) and the first DNA was synthesized from RNA using a PrimeScript RT regent kit (Takara, Shiga, Japan). Actin-coding gene (KP341672) was used as an internal control and the resulting data were normalized using the comparative Cτ method as described previously (106).

### Statistical analysis

The statistical significance of treatments was determined by analysis of variance and means separated by Duncan’s test (*P* < 0.05).

## Acknowledgements

This work was supported by grants from the National Foundation of Natural Science of China (32072362, 32001847, 31571948) and the earmarked fund for China Agriculture Research System (CARS-27) to H.L.

We thank our colleagues in the Institute of Biotechnology, Zhejiang University, for insightful discussions.

